# SUMOylation of the Cardiac Sodium Channel Na_V_1.5 Modifies Inward Current and Cardiac Excitability

**DOI:** 10.1101/2022.12.26.521675

**Authors:** Jin-Young Yoon, Alexander M. Greiner, Julia S. Jacobs, Young-Rae Kim, William Kutschke, Daniel S. Matasic, Ajit Vikram, Ravinder R Gaddam, Haider Mehdi, Kaikobad Irani, Barry London

## Abstract

**Background:** Decreased peak sodium current (I_Na_) and increased late sodium current (I_Na,L_), through the cardiac sodium channel Na_V_1.5 encoded by *SCN5A*, cause arrhythmias. Many Na_V_1.5 post-translational modifications have been reported by us and others. A recent report concluded that acute hypoxia increases I_Na,L_ by increasing a Small Ubiquitin-like MOdifier (SUMOylation) at K442-Na_V_1.5.

**Objective:** To determine whether and by what mechanisms SUMOylation alters I_Na_, I_Na,L_ and cardiac electrophysiology.

**Methods:** SUMOylation of Na_V_1.5 was detected by immunoprecipitation and immunoblotting. I_Na_ was measured by patch clamp with/without SUMO1 overexpression in HEK293 cells expressing wild type (WT) or K442R-Na_V_1.5 and in neonatal rat cardiac myocytes (NRCMs). SUMOylation effects were studied *in vivo* by electrocardiograms and ambulatory telemetry using Scn5a heterozygous knockout (SCN5A^+/-^) mice and the de-SUMOylating protein SENP2 (AAV9-SENP2) or the SUMOylation inhibitor anacardic acid. Na_V_1.5 trafficking was detected by immunofluorescence.

**Results:** Na_V_1.5 was SUMOylated in HEK293 cells, NRCMs and human heart tissue. HyperSUMOylation at Na_V_1.5-K442 increased I_Na_ in NRCMs and in HEK cells overexpressing WT but not K442R-Na_v_1.5. SUMOylation did not alter other channel properties including I_Na,L_. AAV9-SENP2 or anacardic acid treatment of SCN5A^+/-^ mice decreased I_Na_, prolonged QRS duration, and produced heart block and ventricular arrhythmias. SUMO1 overexpression enhanced membrane localization of Na_V_1.5.

**Conclusion:** SUMOylation of K442-Na_v_1.5 increases peak I_Na_ without changing I_Na,L_, at least in part by altering membrane abundance. Our findings do not support SUMOylation as a mechanism for changes in I_Na,L_. Na_v_1.5 SUMOylation may modify arrhythmic risk in disease states and represents a potential target for pharmacological manipulation.

## Introduction

The cardiac Na^+^ channel, Na_V_1.5 (encoded by *SCN5A*), conducts an inward depolarizing Na^+^ current (I_Na_) and controls cell excitability^1^. A wide array of cellular mechanisms modulate I_Na_ under physiological conditions, and abnormal channel expression and function have been implicated in both inherited and acquired arrhythmias. Na_V_1.5 is composed of 4 homologous domains connected by intracellular linkers. The linkers connecting domains I-II and III-IV play important roles in channel function and are key sites of regulation by both binding proteins and post-translational modifications. Post-transcriptional modulators of Na_V_1.5 (microRNAs miR-219, miR-192-5p, and miR-24) alter RNA abundance and translation ^2^. Beta subunits (e.g. Na_V_β1), binding proteins (e.g. FGF12, MOG1, ACTN2), and post-translational modifications (PTMs, e.g. PKA and PKC-mediated phosphorylation, glycosylation, nitrosylation, ubiquitination, acetylation and SUMOylation) modulate channel properties and membrane trafficking ^3,4–6^. Of note, the Na_V_1.5 I–II cytoplasmic linker loop contains the site for phosphorylation by CaMKII^7^ along with a SUMOylation consensus site ^8^.

SUMOylation is a reversible post-translational modification in which proteins termed small ubiquitin-like modifiers (SUMOs) are covalently linked to lysine residues of target proteins^8^. SUMO is conjugated to its substrates through three sequential enzymatic steps: activation, involving the E1 enzyme AOS1/UBA2; conjugation, involving the E2 enzyme UBC9; and substrate modification, through the cooperation of the protein ligases E2 and E3. De-conjugation (de-SUMOylation) occurs through enzymes called SENPs, which recycle free SUMO and unmodify target proteins. In higher order vertebrates, five SUMO paralogs (SUMO1-5) and six SENP family members (1-3, 5-7) have been identified to date.

Functionally, SUMOylation has been shown to be involved in transcriptional silencing ^9, 10^, genomic stabilization ^10^, and the stress response ^11, 12^, although our understanding of tissue-specific functions of the SUMO system is still incomplete. In the heart, SUMOylation is known to play a role modulating contractility and fibrosis. SUMO1 is conjugated to the sarcoplasmic reticulum calcium ATPase 2a (SERCA2a), enhancing its function and preserving its stability ^13^. Of note, SUMO1 is downregulated in heart failure, potentially contributing to contractile dysfunction and suggesting that SUMO1 may hold a therapeutic role.

SUMOylation is also known to play important both direct and indirect roles in ion channel regulation. SUMOylation modifies the stability of CFTR chloride channels, which are mutated in cystic fibrosis ^14^. SUMOylation of the axonal collapsin response mediator protein 2 (CRMP2) controls trafficking and activity of the voltage-gated Na^+^ channel Na_V_1.7 ^15^. In mouse neurons, SUMOylation of the hyperpolarizing-activated cyclic-nucleotide-gated cation channel HCN2 increases its surface expression and the I_h_ current ^16^. In the mouse brain, hyper-SUMOylation of K_V_7.1, K_V_7.2, and K_V_7.3 (via SENP2 deficiency) causes seizures, sinus bradycardia, and heart block, establishing a mouse model for sudden unexplained death in epilepsy (SUDEP) ^17^. Notably, cardiac-specific loss of SENP2 did not cause bradyarrhythmias. Recently, it has been reported that Na_V_1.5 SUMOylation on lysine 442 increases I_Na,L_ in response to acute hypoxia *in vitro* ^6^.

A direct role for SUMOylation/de-SUMOylation in modulating cardiac ion channels and arrhythmic risk *in vivo* has not been reported. Here, we examined the effects of SUMOylation on Na_V_1.5 and I_Na_, and whether changes in Na_V_1.5 SUMOylation modify arrhythmic risk *in vivo*.

## MATERIALS AND METHODS

For detailed methods, see the Online Supplementary Materials. All animal experiments were approved by the Institutional Animal Care and Use Committee (IACUC) at the University of Iowa.

## RESULTS

### SCN5A is SUMOylated

To confirm Na_V_1.5 SUMOylation in our system, anti-SUMO1 antibodies were used to immunoprecipitate exogenous Nav1.5 in HEK293T cells (expressed via transfection) or endogenous Na_V_1.5 in NRCMs. In both cell types, anti-SUMO1 antibodies pulled down Na_V_1.5 (Figure 1A). Further, antibodies to SUMO1 immunoprecipitated Na_V_1.5 in ventricular tissue from failing human hearts explanted at the time of transplantation (Figure 1A).

**Figure 1.**
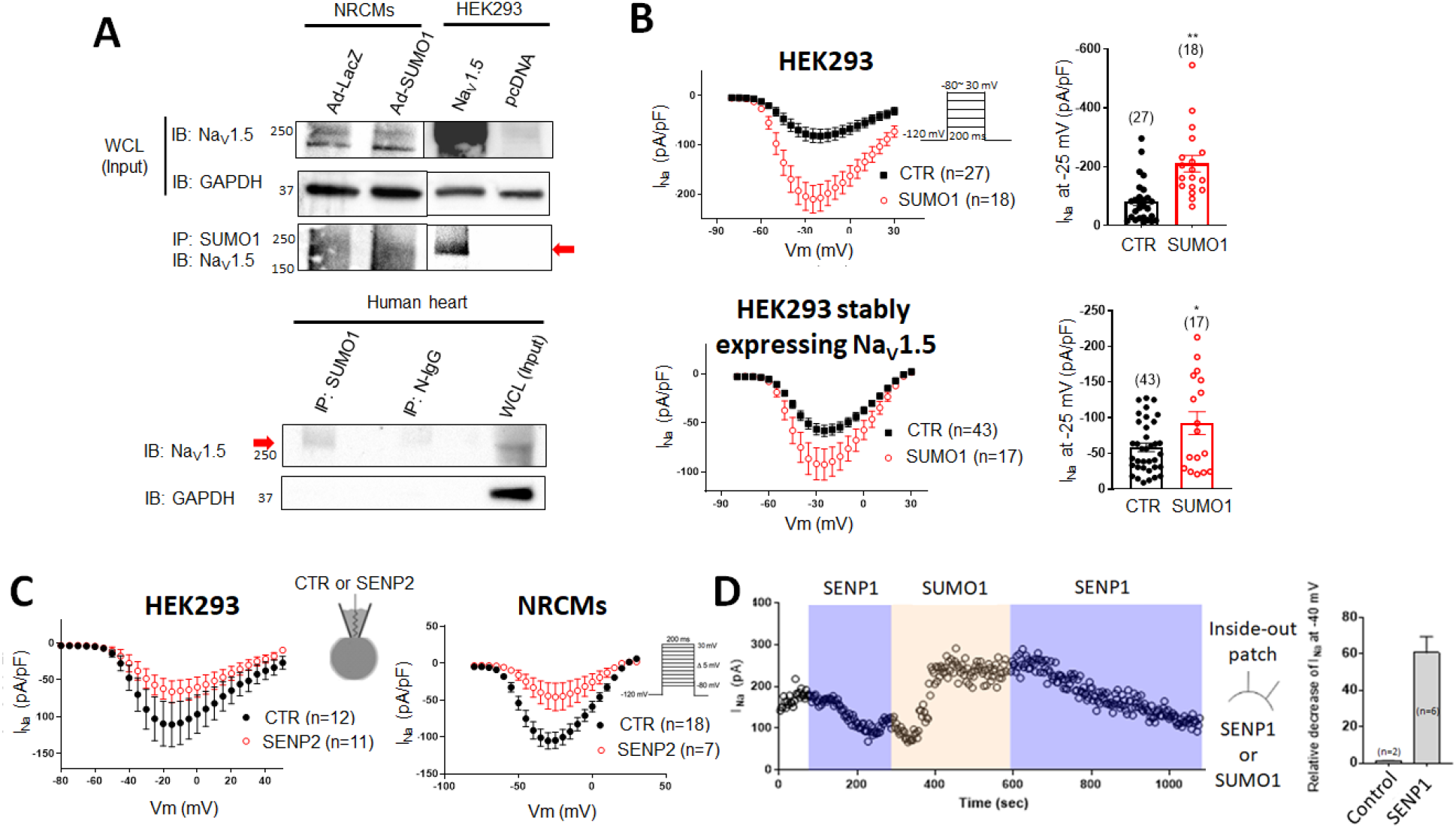
SUMO1 associates with Na_V_1.5 and modulates I_Na_. **(A)** SUMOylation of Na_V_1.5 (arrow) was detected by immunoprecipitation and immunoblotting in NRCMs infected with Ad-SUMO1 and in HEK293 cells transfected with Na_V_1.5 (*upper*) and in ventricular tissue of failing human hearts (*lower*). Adenovirus expressing LacZ (Ad-Lacz) and immunoprecipitation with IgG (N-IgG) were used as controls. **(B)** SUMO1 increases peak I_Na_. Current-voltage (I-V) curves evoked by voltage steps (*inset*) and peak current density (pA/pF) at −25 mV in HEK293 cells transiently co-transfected with Na_V_1.5 (*upper*) or HEK293 cells stably expressing Na_V_1.5 (*lower*) transfected with empty vector (CTR) or SUMO1. (*p < 0.05, **p<0.01). **(C)** Direct application of SENP2 (500 nM) via the patch pipet decreases peak I_Na_ in HEK293 cells expressing Na_V_1.5 (*left*) and NRCMs (*right*). **(D)** SUMOylation and de-SUMOylation reversibly modulate peak I_Na_ in excised inside-out patches from HEK293 cells expressing Na_V_1.5. *Left*, I_Na_ remained constant (192 ± 33 pA) until 100 nM recombinant SENP1 was applied whereupon I_Na_ decreased (80 ± 33 pA). After washout of SENP1, subsequent application of 20 μM recombinant SUMO1 rescued I_Na_, and reapplication of SENP1 gradually reversed the effects of SUMO1. *Right*, Summary data showing that 100 nM SENP1 decreased I_Na_ by ~61% (n=6), whereas current was unchanged in control conditions (n=2).

### SUMOylation increases peak ĪNa

We next examined the effect of SUMOylation on I_Na_ conducted by Na_V_1.5. Using whole cell patch clamp, overexpression of SUMO1 increased peak I_Na_ amplitude in HEK293 cells either transiently transfected with or constitutively expressing Na_V_1.5 (Figure 1B). Direct application of deSUMOylating SENP2 protein through the patch-pipet decreased peak I_Na_ in HEK cells constitutively expressing Na_V_1.5 or in NRCMs expressing endogenous rat Na_V_1.5 (Figure 1C). To verify the effect of SUMOylation and de-SUMOylation in the same cell, we performed an inside-out patch clamp experiment using HEK293 cells overexpressing Na_V_1.5, where the cytoplasmic face of channels was directly exposed to SUMO1 or SENP1. Application of 100 nM SENP1 decreased I_Na_ at −40 mV by ~80%. After removal of SENP1, subsequent exposure to 1μM SUMO1 increased I_Na_, which overshot the baseline levels. Reapplication of SENP1 again decreased I_Na_ (Figure 1D).

### SUMOylation does not increase I_Na,L_

Acute hypoxic conditions were recently reported to SUMOylate Na_V_1.5-K442 and increase late Na_V_1.5 currents (I_Na,L_)^6^. In contrast, SUMO1 overexpression in HEK293 cells or NRCMs did not change I_Na,L_ or the inactivation kinetics of Na_V_1.5 currents (Figure 2A-C). To verify that the observed I_Na,L_ was not contaminated by leak currents, we also measured the tetrodotoxin (TTX, a specific sodium channel blocker)-sensitive current ^18^ in NRCMs. SUMOylation of endogenous Na_V_1.5 in NRCMs did not increase TTX-sensitive I_Na,L_ (Figure 2D). Given that the Na_V_β1 beta subunit can affect I_Na,L_ and inactivation kinetics of Na_V_1.5 ^19, 20^, we co-expressed Na_V_1.5 with the Na_V_β1 subunit in HEK293 cells and showed that SUMOylation did not increase I_Na,L_ in the presence of Na_V_β1 (Figure 2E).

**Figure 2.**
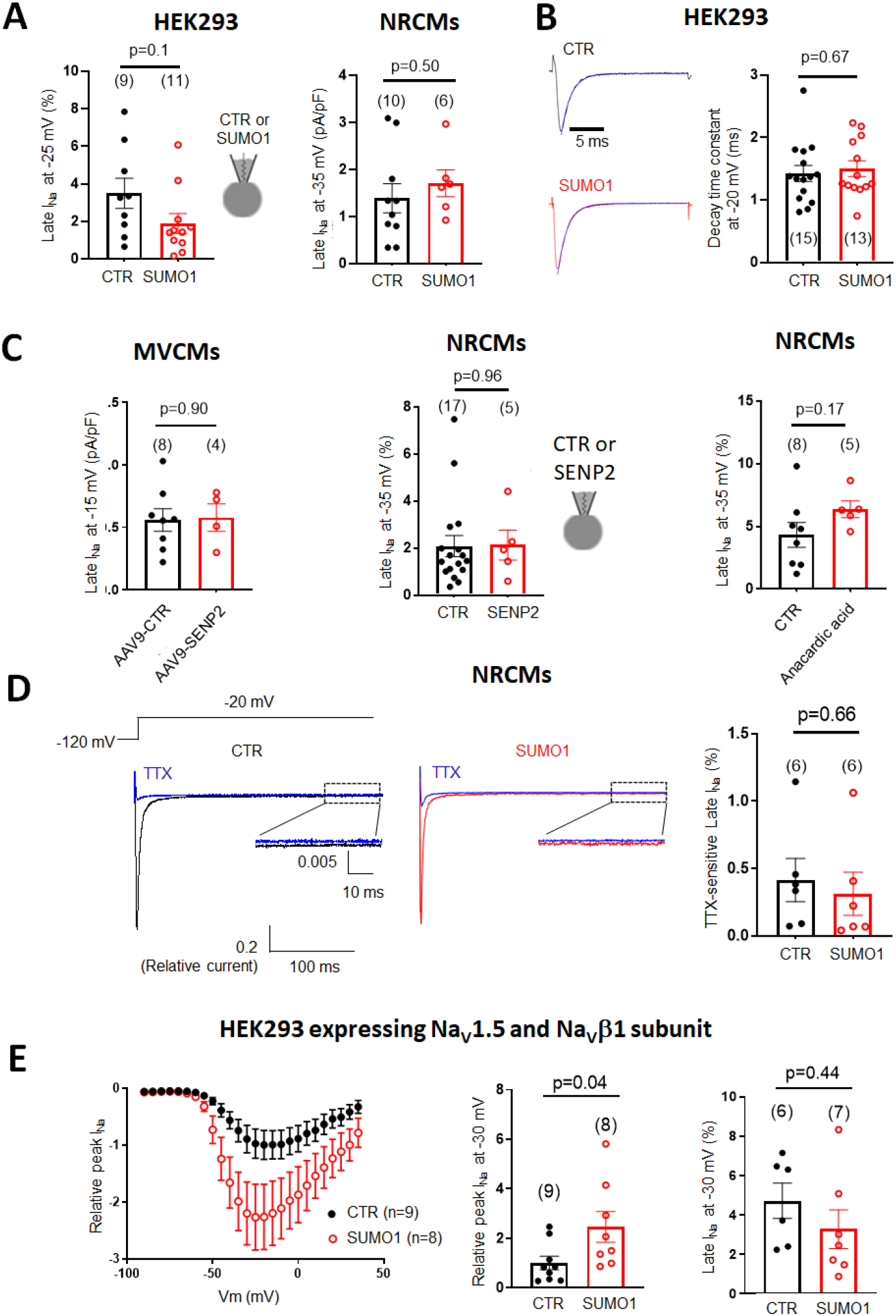
SUMOylation/DeSUMOylation does not change late Na_V_1.5 current or inactivation kinetics. **(A)** ***Left***, I_Na,L_ of HEK293 cells expressing Na_V_1.5 with 20 μM SUMO1 or heat-inactivated SUMO1 protein (CTR) in the patch pipet. ***Right***, I_Na,L_ of NRCMs transfected with either empty vector (CTR) or SUMO1. **(B) *Left*,** Representative Na^+^ currents of HEK293 cells constitutively expressing Na_V_1.5 transfected with either empty vector (black) or SUMO1 (red). Traces were normalized to the peak I_Na_. The decay of the current traces was fit with a mono-exponential function (blue line). ***Right*,** Average time constants for the decay of I_Na_ for control and SUMO1-expressing cells. **(C) *Left*,** I_Na,L_ of MVCMs from the mice injected with AAV9-control virus or AAV9-SENP2 virus. ***Middle***, I_Na,L_ of NRCMs treated with SENP2 (500 nM) or heat-inactivated SENP2 protein (CTR) via the patch pipet. ***Right***, I_Na,L_ of NRCMs treated with vehicle (CTR) or anacardic acid (10 μM, 6hr). **(D)** Quantification of tetrodotoxin (TTX, 30 μM)-sensitive I_Na,L_. I_Na,L_ was measured following a 300 ms voltage steps to −20 mV from a holding potential of −120 mV (***inset***). Average current between 250 and 300 ms was normalized to peak I_Na_. **(E)** I-V curves, peak I_Na_ (normalized to CTR), and I_Na,L_ (normalized to peak I_Na_) of HEK293 cells co-transfected with Na_V_1.5 and Na_V_β1, and either empty (CTR) or SUMO1 vectors.

SUMO1 overexpression caused a subtle left shift of steady state activation (SSA) in HEK293 cells but not in NRCMs, while direct application of SENP2 protein caused a small but statistically significant decrease in the slope of steady state inactivation in both HEK293 cells and NRCMs (Supplemental Figure 1). In addition, SUMO1 transfection caused a small but statistically significant speeding of the fast component of recovery from slow-inactivation in HEK293 cells (Supplemental Figure 2).

### SUMOylation modulates Na_V_1.5 at K442

To identify the SUMOylated lysine residue(s) in Na_V_1.5 modified by SUMO1, we used the SUMOylation consensus sequence (ΨKXD/E, where Ψ is a hydrophobic amino acid, and X is any amino acid), and identified three potential SUMOylation sites in the human Na_V_1.5 α-subunit at K442 (domain I-II linker, intracellular), K1120 (domain II-III linker, intracellular), and K1683 (domain IV pore, extracellular) from among the 83 lysine residues. All of these sites are conserved in mouse, rat, and pig. We focused on K442, which is located in the I-II inter-domain linker (Na_V_1.5 [I-II]) that plays an important role in regulating channel localization and kinetics.^21^

We changed lysine 442 to arginine to prevent SUMO conjugation without disrupting charge interactions (Na_V_1.5-K442R). In transiently transfected HEK293 cells, Na_V_1.5-K442R protein expression and peak I_Na_ were unchanged compared to WT (Figure 3A). Cells expressing Na_V_1.5-K442R showed a small decrease in the slope of steady-state activation compared to WT, with no change in steady-state inactivation (Supplemental Figure 3). Of note, SUMO1 overexpression and direct application of SUMO1 protein increased peak I_Na_ in HEK293 cells expressing WT Na_V_1.5 but not in cells expressing K442R-Na_V_1.5 (Figures 3B, C). In addition, the left shift in SSA seen in WT cells treated with SUMO1 was not seen in cells expressing K442R-Na_V_1.5 (Supplemental Figure 3). Further, we showed that a FLAG-tagged N-terminal fragment of Na_V_1.5 (2.3 kB; carrying the domain I-II linker) co-expressed with SUMO1 requires an intact lysine at K442 to co-precipitate using antibodies against SUMO1 and FLAG (Figure 3D). Together with the patch clamp data, these findings show that K442 is the key Na_V_1.5 residue that regulates I_Na_ by SUMOylation.

**Figure 3.**
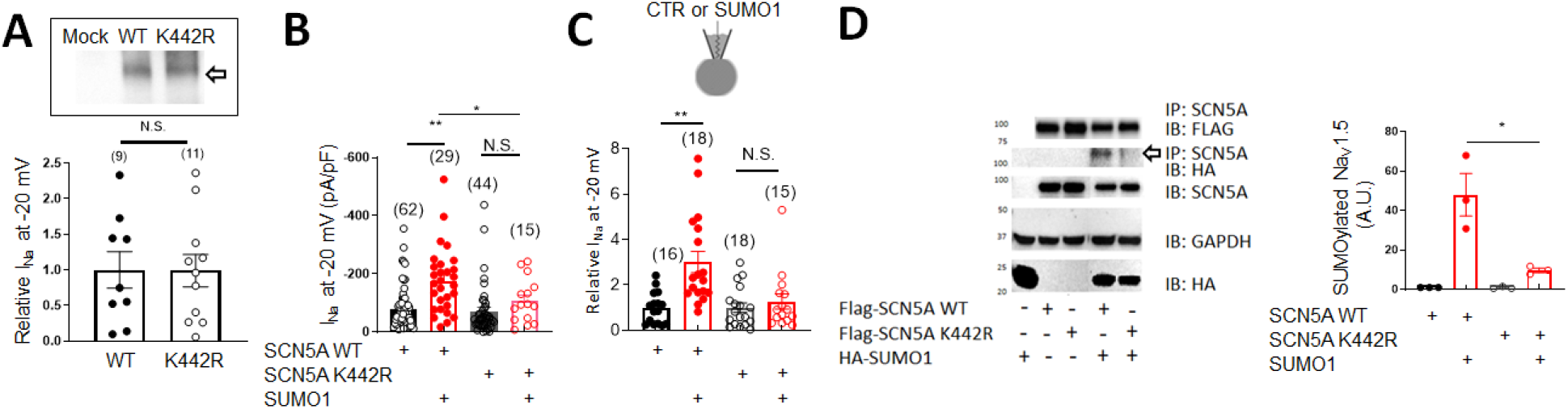
SUMO1 does not SUMOylate or increases I_Na_ in HEK293 cells expressing K442R-Na_V_1.5. (**A**) I_Na_ at −20 mV in HEK293 cells expressing WT- or K442R-Na_V_1.5. I_Na_ was normalized to WT Na_V_1.5 experiments performed from the same batch. ***Insert***, Western blot showing WT- and K442R-Na_V_1.5 expression. SUMO1 overexpression (**B**) or direct application of SUMO1 via patch pipet (**C**) increases I_Na_ in cells transfected with WT-but not with K442R-Na_V_1.5. (**D**) K442R-Na_V_1.5 is not SUMOylated HEK293 cells. *Left*, FLAG-tagged WT- or mutant K442R-Na_V_1.5f N-terminal fragments were expressed in HEK293 cells with/without HA-SUMO1 and immunoprecipitated with an anti-Na_V_1.5 antibody. WT-Na_V_1.5f was SUMOylated (*arrow*) while K442R-Na_V_1.5f was not. *Right*, Quantification of SUMOylated Na_V_1.5 (*n*=3 for each group). *p < 0.05, **p<0.01.

### HypoSUMOylation of Na_V_1.5 evokes an arrhythmic phenotype

Inherited human SCN5A channelopathies are associated with a wide range of cardiac conduction abnormalities and arrhythmic phenotypes. SCN5A mutations that decrease I_Na_ can result in Brugada syndrome and progressive cardiac conduction system defects with bradycardia^22^. In addition, reduced expression of Na_V_1.5 in SCN5A^+/-^ mice leads to conduction delay, atrial and ventricular arrhythmias ^23^. Because diminished SUMOylation activity or expression decreases INa, we looked for cardiac conduction defects and arrhythmias in hypoSUMOylated mouse models.

We first tested the SUMO E1 ligase inhibitor anacardic acid ^24^. Anacardic acid (10 μM) decreased I_Na_ in HEK293 cells expressing Na_V_1.5 (Figure 4A). We then treated a cohort of 7-month-old mice heterozygous for a targeted deletion of Scn5a (Scn5a^+/-^ mice) with 20 mg/kg anacardic acid IP daily for 10 days. High resolution EKGs of treated mice showed mild PR and marked QRS prolongation (Figure 4B, C), indicating that pharmacologic inhibition of SUMOylation impairs cardiac conduction^25^.

**Figure 4.**
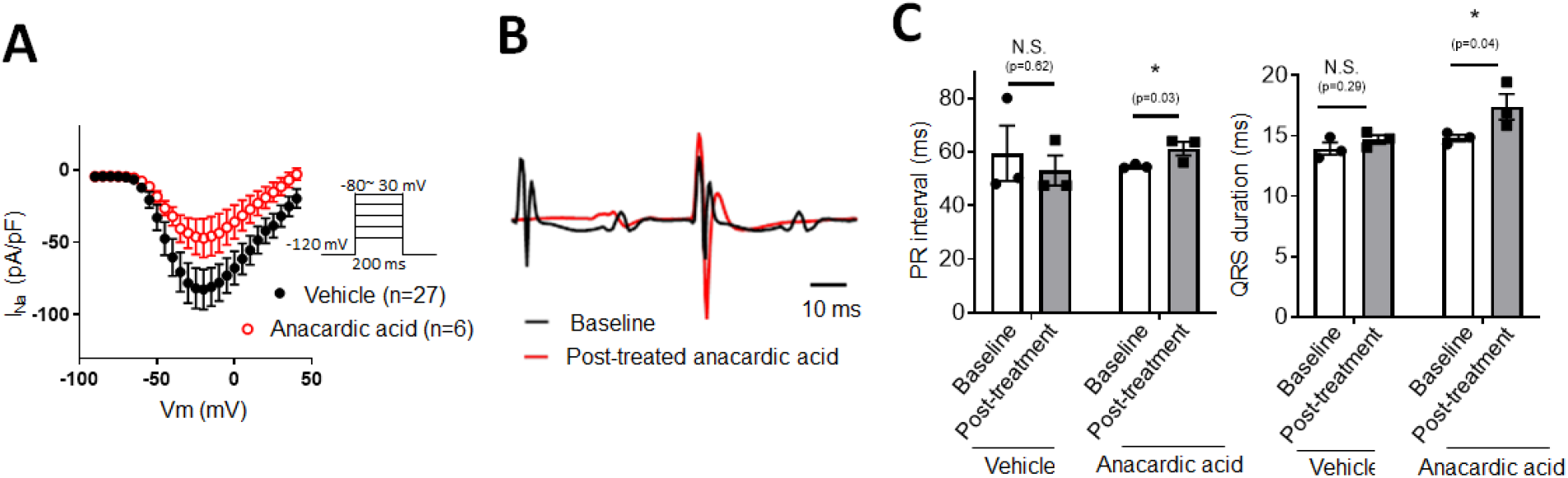
Prolonged PR interval and QRS duration in Scn5a^+/-^ mice dosed with anacardic acid. **(A) 1**0 μM anacardic acid decreases I_Na_ in HEK293 cells expressing Na_V_1.5. Representative ECG traces **(B)** and summary data **(C)** showing mild PR and markedly QRS prolongation in Scn5a^+/-^ mice administered vehicle (n=3) or 20 mg/kg anacardic acid (n=4) for 10 days. *p<0.05.

We then engineered an AAV9-FLAG-SENP2 virus with an IRES-eGFP to more specifically de-SUMOylate proteins in cardiac myocytes. We chose SENP2 because it is expressed in the heart and its main enzymatic activity is de-SUMOylation, as opposed to SENP1 which both promotes and reverses SUMOylation^8^. NRCMs infected with AAV9-FLAG-SENP2 expressed SENP2 mRNA and protein (Figures 5A-C), and had decreased global SUMOylation (Supplemental Figure 5A). Whole-cell patch clamp showed that I_Na_ was significantly decreased in cells expressing FLAG-SENP2 (Figure 5D).

**Figure 5.**
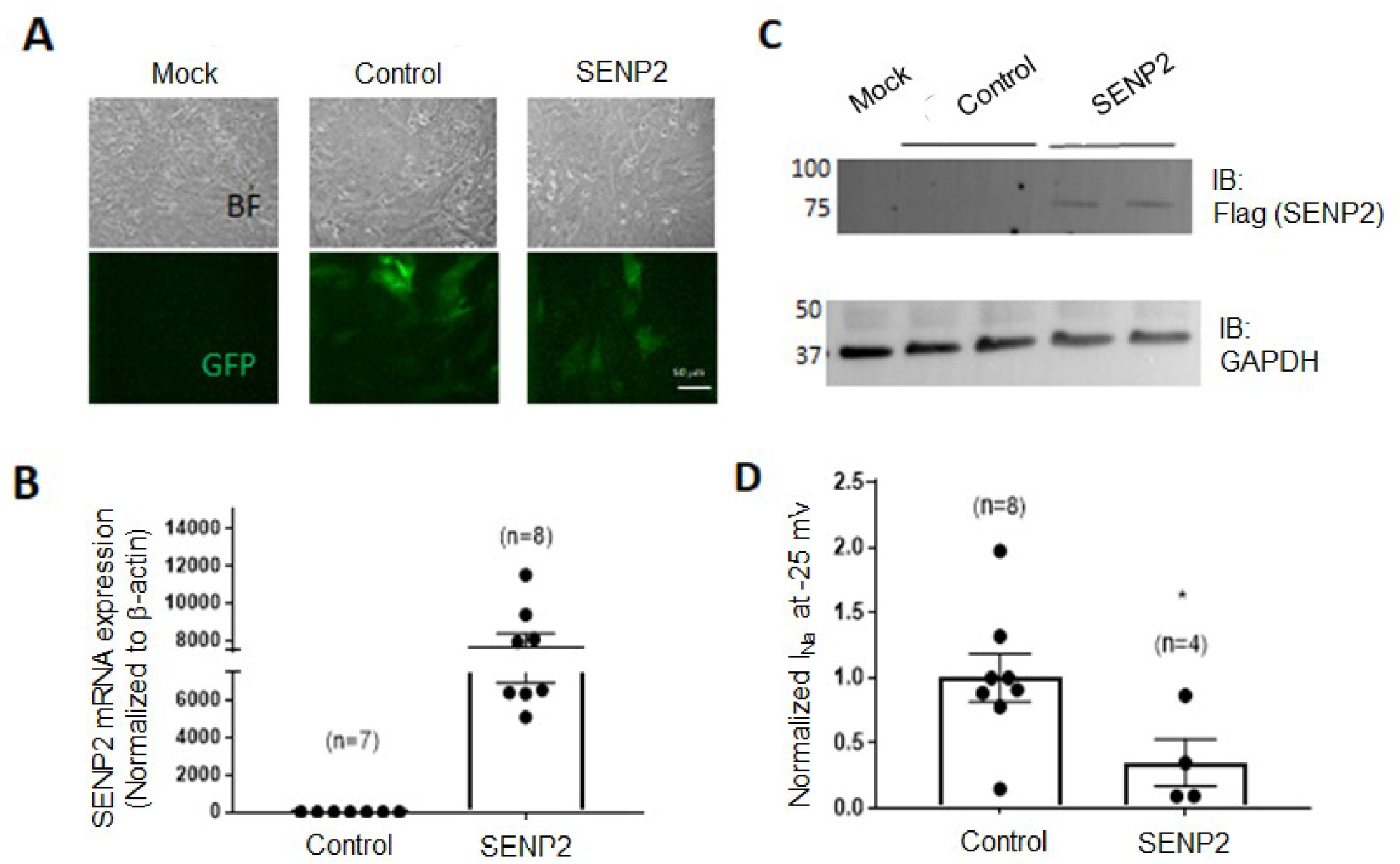
AAV9-SENP2 infection decreases I_Na_ in NRCMs. (**A**) Photomicrographs showing GFP expression in NRCMs infected with control AAV9-eGFP or AAV9-FLAG-SENP2-IRES-eGFP (magnification 20x). AAV9-SENP2 mRNA (**B**) and protein expression (**C**) in NRCMs infected with control or FLAG-tagged AAV9-SENP2. **(D**) I_Na_ at −25 mV is markedly decreased in cells infected with SENP2 compared to control virus. *p<0.05.

We next injected 6-month-old Scn5a^+/-^ mice with AAV9-FLAG-SENP2 IRES-eGFP virus or control AAV9 IRES-eGFP. Hearts from mice injected with both AAV constructs expressed eGFP while only hearts injected with AAV9-FLAG-SENP2 IRES-eGFP expressed FLAG-tagged SENP2 (Figure 6A). High resolution EKGs showed marked prolongation of the QRS duration as early as two weeks following injection with AAV9-FLAG-SENP2 IRES-eGFP (p<0.01, Figures 6B, 6C), with no change in QT, QTc, or PR intervals (Supplemental Figure 4) compared to mice injected with control virus. Radiotelemetry performed four weeks after injection showed bradyarrhythmias including prolonged pauses and high degree AV block as well as ventricular arrhythmias with PVCs and bigeminy in the SENP2 expressing mice but not in controls (Figures 6D-F).

**Figure 6.**
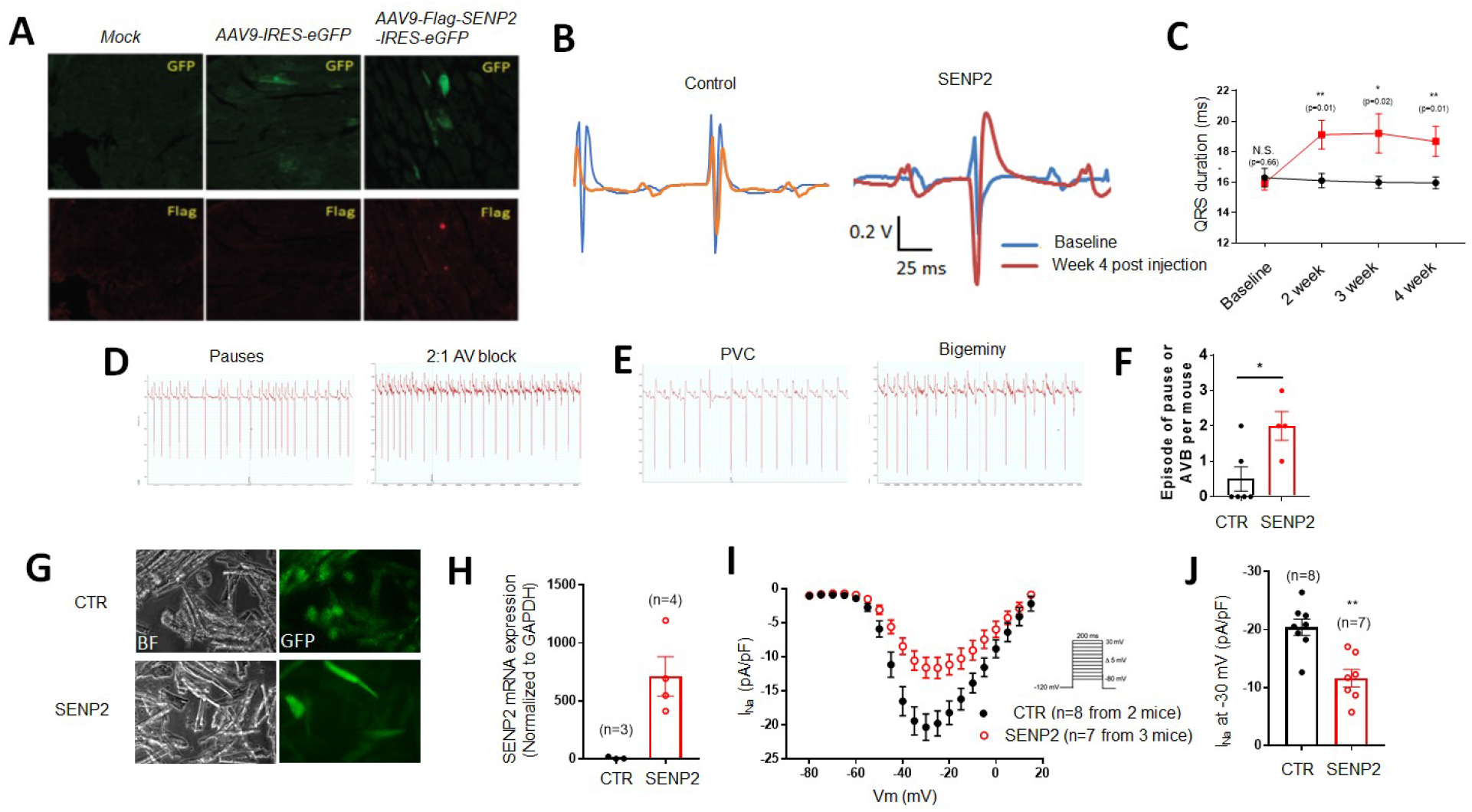
AAV9-SENP2 causes conduction disease and arrhythmias in 6-month-old Scn5a^+/-^ mice. (**A**) Immunohistochemistry showing GFP expression (top) and FLAG expression (bottom) in mice injected with AAV9-IRES-eGFP or AAV9-FLAG-SENP2-IRES-eGFP. Representative EKG tracings **(B)** and summary data **(C)** showing QRS prolongation as early as 2 weeks after infection with AAV9-SENP2. Representative bradyarrhythmias **(D)** and ventricular ectopy **(E)** recorded by ambulatory telemetry 4-weeks after AAV-SENP2 injection. **(F)** Quantification of pauses or atrial-ventricular block (AVB) from the mice shown in ***D-E*. (G)** eGFP expression in MCVMs from mice injected with control (CTR) and AAV9-SENP2 virus. **(H)** Quantitative PCR showing viral SENP2 mRNA expression in MCVMs. **(I, J)** I-V curves and peak I_Na_ measured at −30 mV in MCVMs from mice injected with CTR or AAV9-SENP2 virus. *Inset*, pulse protocol. **p<0.01.

We enzymatically isolated ventricular myocytes (MVCMs) from the hearts of AAV9-SENP2 and AAV9-GFP infected mice. Infection with the viral vectors had no significant effect on cardiomyocyte shape, adherence, or viability. Over 25% of myocytes from mice injected with AAV9-SENP2 expressed eGFP, and SENP2 mRNA was easily detectable by qPCR (Figures 6G, 6H). Global SUMOylation was reduced in MVCMs isolated from AAV9-SENP2 injected versus AAV9-GFP injected controls (Supplemental Figure 5B, 5C). Patch clamp studies showed a marked decrease in I_Na_ in MVCMs expressing GFP from AAV9-SENP2 infected mice (Figures 6I, 6J). There was no change in steady state inactivation, the time constants of the decay phase of I_Na_, (data not shown) or I_Na,L_ (Figure 2C) between control and AAV9-SENP2 infected MCVMs. Taken together, de-SUMOylation in the heart decreases peak I_Na_ and can cause arrhythmias.

### SUMOylation promotes membrane localization of Na_V_1.5

Given the observations that Na_V_1.5 SUMO1 or SENP2 predominantly affect the amplitude of I_Na_ rather its kinetics and that SUMOylation involves changes in expression and stability as reported in SERCA2a and CFTR chloride channels ^13, 14^, we checked the role of SUMOylation on SCN5A expression. Overexpression of SUMO1 or SENP2 did not change Na_V_1.5 mRNA in HEK293 cells, or Na_V_1.5 protein in HEK293 cells, NRCMs, or MVCMs (Supplemental Figure 6).

SUMOylation can affect channel density at the cell membrane ^26^, and modulates ubiquitination and acetylation (via ubiquitin ligase Nedd4-2 and Sirtuin1, respectively) which can affect trafficking of Na_V_1.5 to the cardiomyocyte membrane ^5, 27^. We therefore explored whether SUMOylation controls the trafficking of Na_V_1.5 to the plasma membrane. Quantitative cell surface fluorescence assay using HEK293 cells expressing Na_V_1.5 tagged with an extracellular FLAG epitope and transfected with constructs overexpressing SUMO1 or SENP2 showed that overexpression of SUMO1 increased cell surface expression of Na_V_1.5, whereas expression of SENP2 did not (Figure 7).

**Figure 7.**
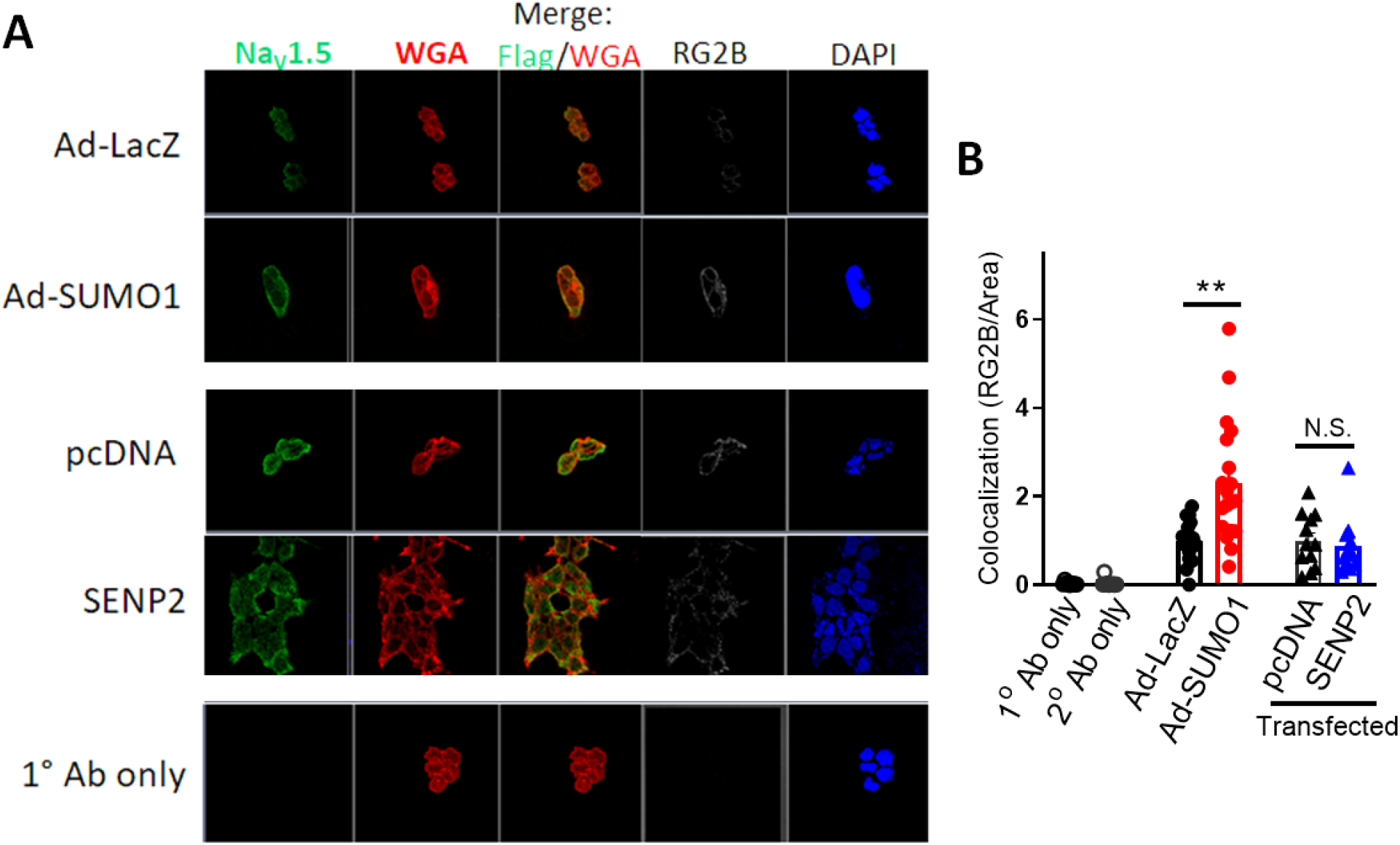
SUMO1 enhances membrane localization of Na_V_1.5 in HEK293 cells. **(A)** immunofluorescence images illustrating colocalization of an extracellular FLAG epitope-tagged Na_V_1.5 (green) and wheat germ agglutinin (WGA, red) in HEK293 cells following adenoviral overexpression of either LacZ (Ad-LacZ), SUMO1 (Ad-SUMO1) or transfection of pcDNA or SENP2. Pixels that colocalize Na_V_1.5 and WGA immunofluorescence were detected using the RG2B co-localization plug-in of Image J. **(B)** Quantitative analysis of the percentage of WGA pixels that colocalize with Na_V_1.5. **p<0.01.

## DISCUSSION

The extent to which different classes of ion channels are SUMOylated and the function and regulation of ion channel SUMOylation are only beginning to be investigated. Na_V_1.5 was previously shown to be SUMOylated at K442 by SUMO1 under acute hypoxic conditions *in vitro* ^6^, associated with increased I_Na,L_ by patch clamp of transfected CHO cells and human pluripotent stem cells induced to a cardiac myocyte phenotype (iPS-CMs). Our study also found that Na_V_1.5 was SUMOylated at K442 in murine ventricular myocytes and human heart samples. In contrast to their work, however, we found that SUMOylation of Na_V_1.5 increased peak I_Na_ amplitude with no change in I_Na,L_, and that deSUMOylation in Scn5a^+/-^ mice was associated with bradyarrhythmias and ventricular ectopy.

### SUMOylation site on Na_V_1.5

The SUMOylated lysine at amino acid 442 sits in the cytoplasmic loop linking domains I and II, a region enriched with arrythmia-causing mutations (at over 30 positions) and previously identified post-translational modifications (see Uniprot). Indeed, this cytoplasmic loop includes five amino acids subject to regulatory phosphorylation. An E446K mutation was reported to cause dilated cardiomyopathy with cardiac arrhythmias, conduction disorder, and decreased I_Na_ ^28, 29^, and an H445D mutation is linked to atrial fibrillation ^30^.

Human Na_V_1.5 contains two other consensus SUMOylation motifs (ΨKX[D/E]), at lysines K496 and K1683, and the three consensus SUMOylation sites are conserved in human, mouse, rat and pig. Our mutational analysis demonstrated that K442 SUMOylation by SUMO1 can regulate I_Na_, and that SUMO1 had no impact on I_Na_ if the site was mutated (K442R). Mass spectrometry analysis in another study only detected SUMOylation of K442 during hypoxia ^6^. Together, these findings suggest that K442 is likely a major site for regulation by SUMOylation. Our findings do not rule out additional regulatory effects by SUMOylation at other lysines or by SUMO2/3. Given that SUMO1 and SUMO2/3 often have different (and sometimes opposite) effects ^8^, future studies would are necessary.

### Role of SUMOylation on Na_V_1.5 channel properties

Plant et al. reported that SUMOylation of K442-Na_V_1.5 in response to acute hypoxia increased I_Na,L_ with no change in peak I_Na_, and that SUMOylation increased the window current. We found an increase in peak I_Na_ with no change in I_Na,L_ or window current. We cannot explain the discrepancy between these studies. We considered a possible role for Na_V_1.5 β-subunits, but co-expression of Na_V_β1 did not change our results. Our finding in cardiac Na_V_1.5 resembles the finding that acute hypoxia mediates SUMOylation of the neuronal sodium channel Na_V_1.2 on the surface of central neurons, leading to an increased peak current without a change in late current ^31^. Additionally, our findings are also consistent with the QRS widening without change in QTc seen in our hypoSUMOylated mouse model treated with SENP2. Potential explanations for the discrepancy could include global or regional changes in oxidative stress caused by hypoxia, differences in our experimental systems, effects of other α and β Na^+^ channel subunits, and/or differences in experimental pulse protocols.

### Indirect modulation of Na_V_1.5 SUMOylation

SUMOylation is known to play both important direct and indirect roles in ion channels and can affect protein-protein interactions ^8^. Although our data show that Na_V_1.5 is a direct target for SUMOylation by SUMO1, global SUMOylation of other proteins might indirectly regulate Na_V_1.5. For example, we have shown that SIRT1 increases I_Na_ by deacetylating Na_V_1.5 ^5^; SIRT1 activity is affected by SUMOylation ^32^. SUMOylation and ubiquitination can occur on the same lysine residues and may compete with each other to alter trafficking of Na_V_1.5 to and from the cardiomyocyte membrane ^5, 27^. Reactive oxygen species (ROS) originating from the mitochondria inhibit peak I_Na_ and increase I_Na,L_, and SUMO proteins can regulate mitochondrial function ^33^. SUMOylation of SERCA2a, which is downregulated in heart failure, enhances its function and preserves its stability ^13^. Diminished SERCA2a activity might increase basal intracellular Ca^2+^ and CaMKII activity ^34^, modifying Na_V_1.5 properties ^21^. Thus, we cannot rule out indirect effects of SUMOylation on Na_V_1.5 under both normal and pathological conditions.

## Conclusions

SUMOylation of Na_V_1.5 at K442 increases peak I_Na_ with no change in I_Na,L_, at least in part through increased membrane trafficking, and SUMOylation deficiency may increase arrhythmia risk in the setting of Na^+^ channel deficiency. These findings provide another potential mechanism underlying the pathogenesis of cardiac arrhythmias. In addition, they provide a rationale for testing SUMO1 activators in pathological conditions characterized by decreased cardiac Na^+^ current.

## Acknowledgements

We thank Kristina Greiner and Diana Colgan in the Carver College of Medicine at the University of Iowa for assistance writing and proofreading.

