## Supplementary material for "SUMOylation of the Cardiac Sodium Channel Na_V_1.5 Modifies Inward Current and Cardiac Excitability": MATERIALS AND METHODS, and Supplemental Figures 1 to 6

### SUPPLEMENTAL INFORMATION

#### MATERIALS AND METHODS

***Animals***. SCN5A$^{+/-}$ mice (backcrossed more than 10 times onto the C56BL6/J strain) were a gift from Dr. Dan M. Roden (Vanderbilt University, Nashville, TN). Mice were housed in a controlled temperature environment on a 12-hour light/dark cycle, with food and water provided ad libitum. All studies were approved by the Institutional Animal Care and Use Committees (IACUC) at the University of Iowa.

***Human heart samples.*** Human myocardial tissue was extracted under protocols approved by Institutional Review Board of the University of Iowa. Ventricular tissue from failing human hearts explanted at the time of transplantation was collected. Hearts were arrested in situ using ice-cold cardioplegia solution and followed immediately by freezing in liquid nitrogen and storage at −80 °C until use.

***Recombinant AAV production and infection in vitro or in vivo.*** AAV9- Flag -SENP2-IRES eGFP was produced using plasmids pFBAAVCMVmcswtIRESeGFPBgHpA by the University of Iowa Viral Vector Core Facility. Six- to 9-month-old male SCN5A$^{+/-}$ mice received AAV9- Flag -SENP2-IRES eGFP or AAV9-eGFP control virus at a dose of 8.6 x $10^{12}$ vg/kg BW via jugular-vein injection. Cardiac function was measured weekly for 6 weeks after infection. NRCMs were infected at 37°C with AAV9- Flag -SENP2-IRES eGFP or AAV9-eGFP control virus at 100 multiplicities of infection, and NRCMs were studied 48 hours later [1].

***Reagents.*** Antibodies were used to Na$_V$1.5 (Abcam [Cambridge, MA]-ab63288, Almone Labs [Jerusalem, Israel]-ASC-005, Millipore [Burlington, MA]-AB5493), Sirt1 (Santa Cruz Biotechnology [Dallas, TX]-sc-15404), and GAPDH (Trevigen [Gaithersburg, MD]-2275-PC).

All other reagents were procured from Sigma-Aldrich (St. Louis, MO) unless otherwise specified. The $Na_V1.5$ mutant constructs were generated by site-directed mutagenesis.

***Cell culture.*** HEK293 cells were obtained from the American Type Culture Collection (ATCC, Manassas, VA) and were cultured in 10% FBS-supplemented DMEM media.

***NRCM preparation and transfection.*** NRCMs were isolated from 1- to 2-day-old Sprague-Dawley rats (Harlan Laboratories, Indianapolis, IN). The hearts were surgically extracted and the ventricles digested overnight using a standardized trypsin and collagenase-based protocol (Worthington Biochemical, Lakewood NJ)[2]. Isolated cells were plated in a NRCM culture medium (a mixture of Dulbecco's modified Eagle's medium [DMEM] with F12 supplemented with HEPES, glutamine, insulin-transferrin-selenium [1], bromodeoxyuridine [to inhibit fibroblast replication], 5% stripped horse serum, and gentamycin [50 µg/ml, Life Technologies, Carlsbad, CA]) on glass coverslips coated with laminin and stored at 37°C in a 5% $CO_2$ incubator. After 2 h following the isolation, cells were transfected with using Lipofectamine 2000 (Life Technologies), according to the manufacturer's protocol. The media was then changed 3 h later to NRCM culture medium (in an equivolume mixture of DMEM and F-12 media containing 5% FBS).

***Immunoprecipitation and immunoblotting.*** Immunoprecipitation of SUMO1 and $Na_V1.5$ channels was carried out by incubating 2 µg of the respective antibodies with 1 mg of cell lysate or tissue homogenate overnight, followed by addition of 50 µl of protein A–Sepharose slurry (Amersham, Piscataway, NJ) for 4 h. After washing, immunoprecipitates were heated to 50°C in SDS–PAGE gel loading buffer, subjected to SDS–PAGE, transferred to PVDF membrane and probed with the specified primary antibody and the appropriate peroxidase-conjugated secondary antibody (GE Healthcare Life sciences, Marlborough, MA). Western blotting of 50 µg of whole-cell lysates (WCL) was similarly performed by using the appropriate primary and secondary antibodies. Chemiluminescent signal was detected using SuperSignal West Pico/Femto substrate (Pierce, Rockford, IL), and blots were imaged with a GelDoc 2000 Chemi Doc system (Bio-Rad, Hercules, CA).

***Determination of cell surface expression of $Na_V1.5$ in HEK293 cells***. A stable HEK293 cell line expressing full-length Nav1.5 tagged with an extracellular FLAG-myc epitope was used[2]. Cells were plated in a 12-well dish at a density of $1.0 \times 10^5$ cells/well and subsequently infected with an adenoviral vectors expressing SUMO1 (Ad-SUMO1) or an Ad-LacZ control vector, or transfected with a SENP2 overexpression plasmid or a pcDNA control plasmid, followed by incubation for 48 h. The cells were then placed on ice to stop additional cellular trafficking. Nonpermeabilized cells were then treated with primary α-FLAG-M2 antibody (1:50 in PBS/0.5% BSA) (Sigma-Aldrich, St. Louis, MO) for 90 min followed by washing with 1×-PBS/0.5% BSA (3×). Cells were then treated with a AlexaFlour488 conjugated-goat anti-mouse secondary antibody (1:1,000) (Invitrogen, Waltham, MA) and co-stained with AlexaFluor594-WGA for 15 min  Cells were washed again with only 1×-PBS and treated with DAPI (0.5 μg/ml) for 1 min followed by a 5-min rinse in 1× PBS. Images were acquired to detect fluorescence of AlexaFluor488, AlexaFluor594, and DAPI along with transmitted light imaging using a Zeiss confocal microscope (Model 710). $Na_V1.5$ pixels that colocalize with WGA were detected using the RG2B co-localization plug in of Image J and normalized with cell surface area.

***Patch-clamp.*** The whole-cell patch-clamp technique was used to measure $Na^+$ current in isolated mouse ventricular myocytes as previously described [2]. Quiescent rod-shaped ventricular myocytes from experimental group mice and age/sex-matched littermate controls were used. For voltage clamp studies, command pulses were generated using pClamp 10 (Molecular Devices,

San Jose, CA) and currents were sampled at 20 KHz through an A/D converter (DigiData 1440, Molecular Devices, CA) and low pass filtered at 5 kHz. An Axopatch 200B patch clamp amplifier (Molecular Devices) was used with ≈ 85% series resistance compensation, yielding a maximum voltage error of ~1mV. The pipette solution consisted of (in mmol/liter) 125 CsCl, 1 $MgCl_2$, 10 EGTA, 5 Na-ATP, and 10HEPES (pH 7.2 with CsOH). The bath solution consisted of (in mmol/liter) 100 TEA chloride, 40 NaCl, 10 glucose, 1 $MgCl_2$, 5 CsCl, 0.1 $CaCl_2$, 1 NiCl, and 10 HEPES (pH 7.4 with CSOH). All experiments were performed at room temperature (20-22 °C). Cellular capacitance was measured, and $I_{Na}$ was normalized to cellular capacitance. Steady state activation and inactivation kinetics were measured. Studies were performed on at least 10 myocytes from at least 3 mice of each genotype.

For recordings from isolated rat neonatal myocytes, the pipette solution contained (in mmol/liter) NaF 10, CsF 120, CsCl 20, EGTA 10, HEPES 10 titrated to pH 7.4 with CsOH and the bath solution contained (in mmol/liter) NaCl 25, NMDG 118.2, KCl 4, $MgCl_2$ 1, $CaCl_2$ 1.8, HEPES 10, glucose 10 and titrated to a pH of 7.4 with HCl. To record $Na^+$ currents from HEK293 cells, electrodes of 1–2 MΩ were filled with a pipette solution containing (in mmol/liter) NaF 10, CsF 110, CsCl 20, EGTA 10 and HEPES 10 (pH 7.35 with CsOH), and the bath solution contained (in mmol/liter) NaCl 40, unless otherwise specified, 103 NMDG, KCl 4.5, $CaCl_2$ 1.5, $MgCl_2$ 1, and HEPES 10 (titrated to pH 7.35 with CsOH). For excised inside-out patches, bath solution (intracellular) contained (in mmol/liter) CsF 40, CsCl 120, EGTA 2, HEPES 10, NaCl 1, $CaCl_2$ 0.68, $MgCl_2$ 2, MgATP 2. The pipet solution (extracellular) contained (in mmol/liter) CsCl 1, HEPES 10, NaCl 160, $CaCl_2$ 2, $MgCl_2$ 1, Glucose 5, MgATP 2. For all solutions osmolality was 300 ± 10 mmol/kg and pH was 7.2. Patches were held at -100 mV, and the currents were evoked from a 50 ms pre-pulse to -120 mV by 20 ms test pulses to -40 mV. Gap-free protocols were utilized.

***Surface electrocardiograms (EKG).*** Mice were anesthetized by isoflurane (1–2%) inhalation and anesthesia was maintained during the procedure by use of a nose cone. High-resolution multilead EKGs were performed using a Data Acquisition System (iWorx/IX-100B, Dover, NH) and subcutaneous bipolar electrodes in positions corresponding to human leads I (right to left shoulder), II (right shoulder to left thigh), III (left shoulder to left thigh) and modified chest V (back to left midclavicular line in the 5th interspace). Electrodes were placed subcutaneously by lightly puncturing through the skin. Signals were digitalized, averaged, and analyzed using LabScribe2 (iWorx) by an investigator blinded to group. Heart rate, and duration of the PR, QRS and QT intervals on the EKG were measured. The QT interval was corrected for cycle length using the variant of Bazzett's formula modified for the mouse [3]. Five to ten consecutive QRS complexes were signal-averaged to improve resolution for the interval measurements.

***Mouse ambulatory telemetry.*** Radiotelemetry electrocardiographic monitors (model ETA-F10, Data Sciences International, New Brighton, MN) were implanted subcutaneously on the abdominal side of mice following isoflurane (2–5%) inhalation for anesthesia. Subcutaneous injection of flunixin meglumine (2.5 mg/kg) was used for analgesia. At least 6 days after the implant, 24-48 hr of telemetry was recorded from the single lead bipolar EKG interfaced with PhysioTel Receiver RPC-1 model (Data Sciences International). Data were sampled at 400 Hz and converted to digital format using the PowerLab data acquisition system and LabChart 8 software (AD Instruments, Colorado Springs, CO). Telemetry data were analyzed for atrial and ventricular arrhythmias by an experienced observer blinded to treatment group.

***Transthoracic echocardiograms.*** Echocardiograms to measure ventricular size, wall thickness, and ejection fraction were performed on mice anesthetized with midazolam using the Vevo 2700 VisualSonics System (Toronto, ON, Canada).

***Sample size and Statistical Analyses.*** Six to 12 mice in each group were used to achieve a power of at least 0.8 for each group with significance defined as $p<0.05$. We performed all statistical analyses using GraphPad Prism (San Diego, CA). To compare all study groups of each experiment on a continuous response variable, we used two-way ANOVA followed by post-hoc analysis using tests appropriate to the data set. Data were analyzed using two-way repeated measures ANOVA. For patch clamp studies, two tailed-unpaired t tests were used to compare experimental and control groups. $p< 0.05$ was considered statistically significant for all tests.

#### SUPPLEMENTAL FIGURES

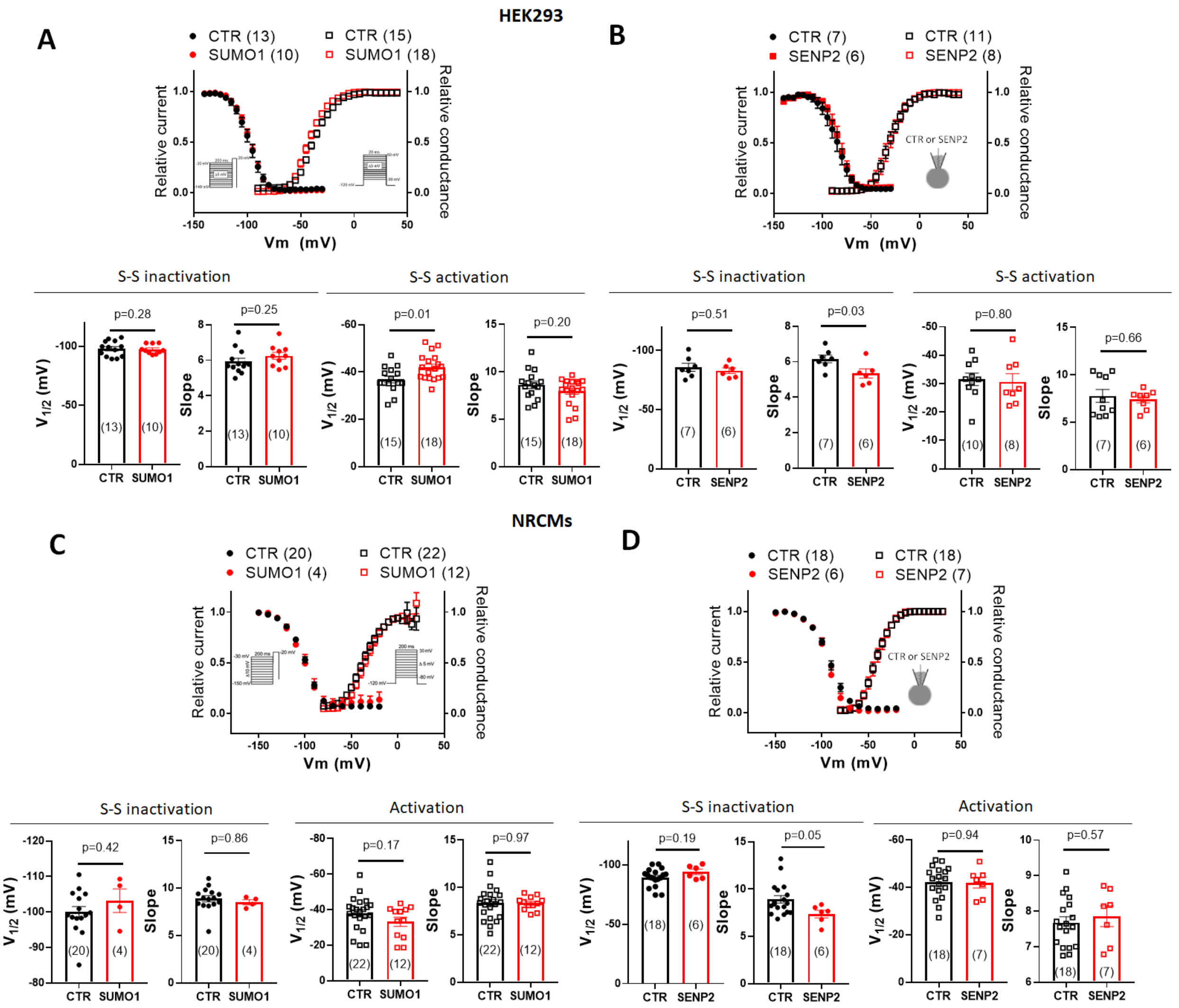


**Supplemental Figure 1.** Effect of SUMOylation/DeSUMOylation on gating properties of steady-state activation and inactivation of Na$_V$1.5. **(A)** Voltage-dependence of steady-state (S-S) inactivation (circles) and activation (squares) of $I_{Na}$ in HEK293 cells expressing Na$_V$1.5 co-transfected with pcDNA (CTR) or SUMO1. S-S inactivation was measured by a two-pulse protocol with 200 ms conditioning pulses from a holding potential of −140 mV to −30 mV in 10 mV increments followed by a 20 ms test pulse to −20 mV (***inset, left***). S-S activation was assessed from a holding potential of -120 mV using a 300 ms test pulses between -90 and 50 mV, in 5 mV increments (***inset, right***). A 5 s interpulse interval was used in both cases. Normalized peak currents were plotted against either prepulse potential or test potential . Normalized activation and inactivation relationships were fit with a Boltzmann function, $I = I_{max}/(1 + \exp[V – V½/k])$, where $I_{max}$ is the maximum current and k is slope factor. Data are mean ± S.E.M. **(B)** SSA and SSI in HEK293 cells transiently transfected with $Na_V1.5$ and treated with 500 nM SENP2 or heat-inactivated protein (CTR) in the pipette. **(C)** Voltage dependence of SSA and SSI of NRCMs transiently-transfected with pcDNA (CTR) or SUMO1; protocols and analysis was similar to (A). **(D)** SSA and SSI in NRCMs treated with 500 nM SENP2 or heat-inactivated protein (CTR) via the pipette. *Inset,* pulse protocol.

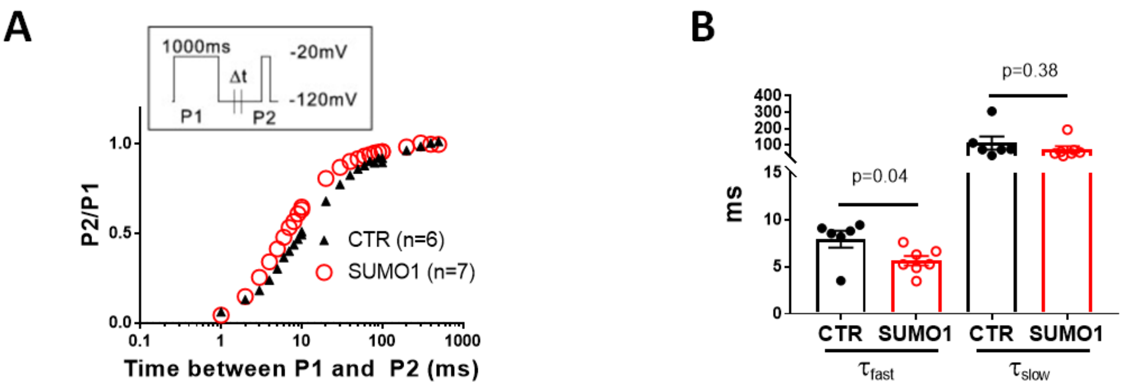


**Supplemental Figure 2. SUMOylation on recovery from inactivation of Nav1.5 channel. (A)** Recovery from slow inactivation assessed after a 1000 ms inactivating prepulse (P1) to −20 mV (P2) (***inset***) in HEK293 cells transiently transfected with $Na_V$1.5 and co-transfected with empty vector (CTR) or SUMO1. **(B)** Time constant for recovery $\tau_{fast}$ and $\tau_{slow}$ were obtained by fitting the normalized current amplitude to the recovery time using the bi-exponential function.

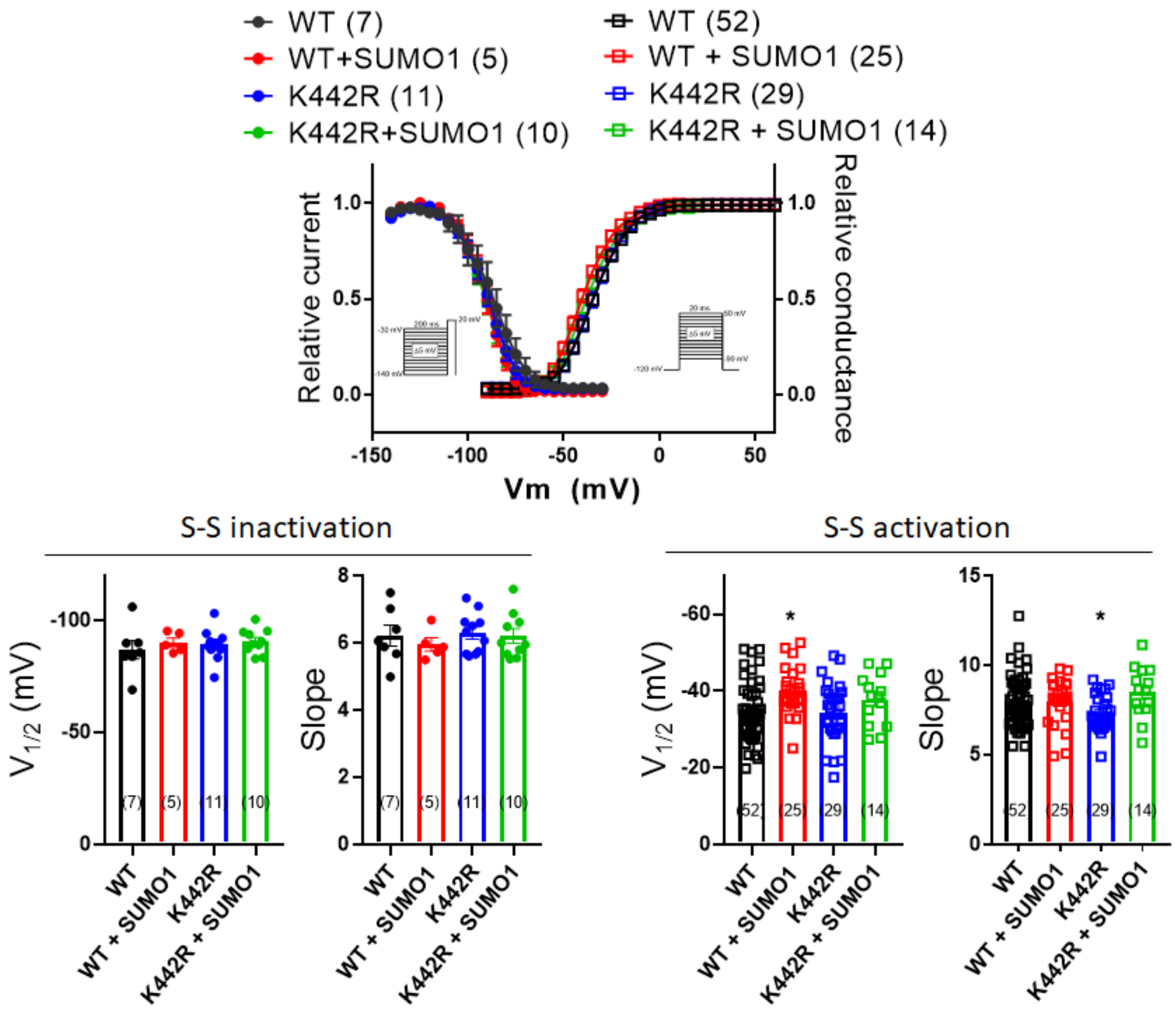


**Supplemental Figure 3.** Comparison of Steady State Activation (SSA) and Steady State Inactivation (SSI) between WT $Na_V1.5$ and $Na_V1.5$- K442R. ***Upper,*** $Na_V1.5$-WT or $Na_V1.5$-K442R were express in HEK293 cells and co-transfected with pcDNA or SUMO1. SSI (circles) and SSA (squares) were measured as described in ***Supplemental Figure1***. ***Lower,*** Summarized values of V½ and slope. Changes from values measured from cells expressing WT-$Na_V1.5$ under control conditions were assessed by ANOVA and are indicated with $*p<0.05$.

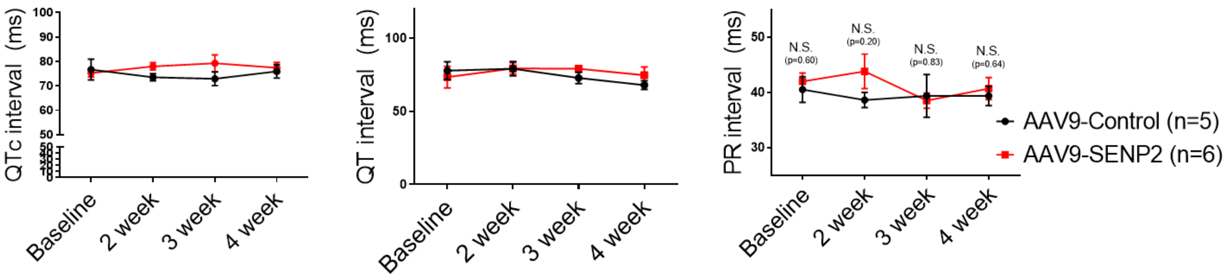


**Supplemental Figure 4.** QT, QTc, and PR intervals from mouse EKG as a function of time after infection with AAV9-SENP2 or control virus.

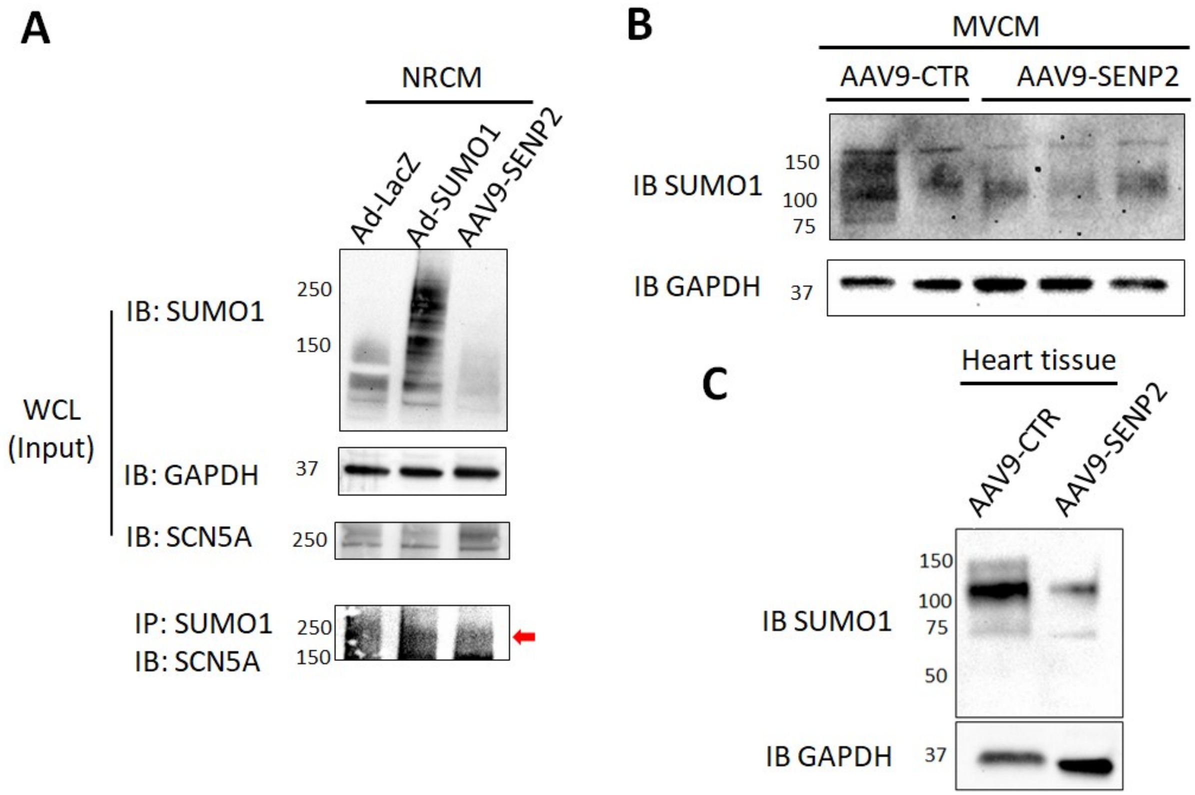


**Supplemental Figure 5. Efficacy of SUMO1 or SENP2 overexpression on either global SUMOylation or SUMOylated-Nav1.5. (A)** NRCMs were infected with either Ad-LacZ, Ad-SUMO1 or AAV9-SENP2. SUMOylation of total protein from isolated mouse cardiomyocytes **(B)** or from mouse heart tissue **(C)**, following injection of AAV9-control or AAV9-SENP2 virus. The heart tissue experiment was performed in duplicate with similar results.

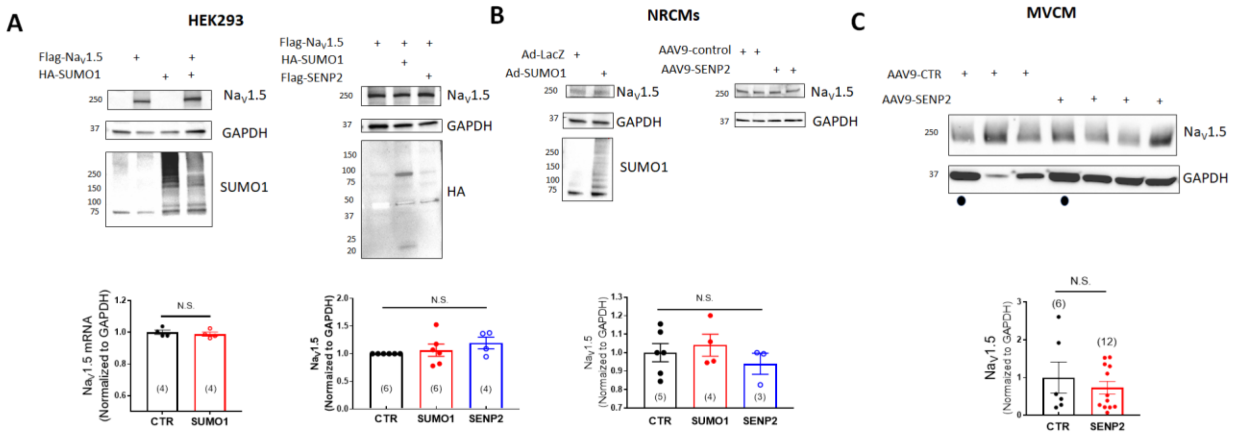


**Supplemental Figure 6. SUMOylation did not affect Nav1.5 expression**. **(A)** Effect of overexpression of SUMO1 or SENP2 on expression of Nav1.5 at mRNA and protein in HEK293 cells transfected with Nav1.5. **(B)** Effect of adenoviral overexpression of SUMO1 or SENP2 on Nav1.5 protein expression in NRCMs. **(C)** Immunoblots showing Nav1.5 expression in MVCM isolated from mice injected with control virus or SENP2 virus. Changes from values measured from cells under control conditions were assessed by ANOVA or two-tailed t-test.
